# Consequences of aneuploidy in human fibroblasts with trisomy 21

**DOI:** 10.1101/2020.08.14.251082

**Authors:** Sunyoung Hwang, Paola Cavaliere, Rui Li, Lihua Julie Zhu, Noah Dephoure, Eduardo M. Torres

## Abstract

An extra copy of chromosome 21 causes Down syndrome, the most common genetic disease in humans. The mechanisms by which the aneuploid status of the cell, independent of the identity of the triplicated genes, contributes to the pathologies associated with this syndrome are not well defined. To characterize aneuploidy driven phenotypes in trisomy 21 cells, we performed global transcriptome, proteome, and phenotypic analysis of primary human fibroblasts from individuals with Patau (trisomy 13), Edwards (trisomy 18), or Down syndromes. On average, mRNA and protein levels show a 1.5 fold increase in all trisomies with a subset of proteins enriched for subunits of macromolecular complexes showing signs of post-transcriptional regulation. Furthermore, we show several aneuploidy-associated phenotypes are present in trisomy 21 cells, including lower viability and an increased dependency on the serine-driven lipid biosynthesis pathway to proliferate. Our studies present a novel paradigm to study how aneuploidy contributes to Down syndrome.

## Introduction

An abnormal number of chromosomes or aneuploidy accounts for most spontaneous abortions as missegregation of a single chromosome during development is often lethal (Nagaoka et al., 2012). Individuals with trisomies of chromosomes 13 or 18, which cause Patau and Edwards syndromes, respectively, are born with severe developmental defects and die soon after birth. Only patients with trisomy 21, which causes Down syndrome can live to adulthood but show developmental abnormalities, cognitive disabilities, congenital heart defects, increased risk for leukemias and neurodegenerative disease, autoimmune disorders, and clinical symptoms associated with premature aging (Hassold and Jacobs, 1984). Importantly, the incidence of aneuploidy is associated with tumorigenesis and increases with age in both somatic and germline tissues in apparently healthy individuals (Holland and Cleveland, 2009; Naylor and van Deursen, 2016). The mechanisms by which aneuploidy affects cellular function to cause Down syndrome, its role in cancer, or how it promotes aging are not well understood.

The phenotypes associated with trisomy 21 at the organismal level are complex as individuals with Down syndrome are born with varying severities of and higher risks for several human diseases (Antonarakis et al., 2020). There are two main gene-centric hypotheses proposed to explain the biological consequences of trisomy 21. One is that increased expression of a particular gene and its downstream pathways cause the pathophysiology in Down syndrome. For example, increased expression of the amyloid-beta precursor protein APP, the High Mobility Group Nucleosome Binding Domain 1 HMGN1, or the transcription factor RUNX1 is associated with the increased incidence of early onset of Alzheimer’s disease, B cell acute lymphoblastic leukemia, or transient myeloproliferative disorder in individuals with Down syndrome, respectively (Elagib et al., 2003; Goldgaber et al., 1987; Lane et al., 2014). A second hypothesis is that abnormalities arise due to the indirect effect of increased gene activity on chromosome 21 by changing the physiology of the cell and altering homeostasis. Examples include genes that regulate splicing (U2AF1L5, RBM1, U2AF1), chromatin regulators (HMGN1, BRWD1), secretory-endosomal functions (DOPEY2, CSTB, and SYNJ1), protein turnover (USP25), or metabolism (SOD1). Nonetheless, it has been challenging to prove that a third wild type copy of a given gene encoded on chromosome 21 is solely the main driver for a particular disease (Antonarakis et al., 2004; Antonarakis et al., 2020).

A third less well-explored consequence of trisomy 21 is the disruption of cellular physiology due to the presence of an extra chromosome independent of the identity of the genes encoded on it; that is, the aneuploid status of the cell. Several studies have revealed aneuploidy-driven phenotypes shared among different organisms and independent of the abnormal karyotype (Santaguida and Amon, 2015; Torres et al., 2008). These include defects in cell proliferation, signs of proteotoxic stress, genomic instability, and altered metabolism. Recently, we showed that in both yeast and human cells, the morphology of the nucleus is significantly affected by the presence of an extra copy of a chromosome (Hwang et al., 2019). This phenotype is associated with dysregulation of the synthesis of lipids that constitute the nuclear membrane. In particular, we found that increasing the levels of long-chain bases suppresses nuclear abnormalities and improves the fitness of aneuploid cells, including human fibroblasts with trisomy 21 (Hwang et al., 2017; Hwang et al., 2019).

To further characterize the phenotypes associated with aneuploidy in human cells, we perform a global transcriptome and proteome quantification of primary fibroblasts isolated from individuals with Down, Edwards, or Patau syndrome. We also found that several aneuploidy-associated phenotypes are present in primary fibroblasts with trisomy 21. These include lower viability, altered lipid metabolism, and an increased dependency on serine to proliferate. Altogether our studies indicate that in addition to the gene-driven pathologies associated with Down syndrome, cellular defects associated with aneuploidy independent of the identity of the genes encoded on chromosome 21 can contribute to disease.

## Results

### Transcriptome analyses of primary fibroblasts show a proportional relationship between copy number and transcript levels

Several studies have suggested a complicated relationship between copy number, mRNA, and protein levels of genes encoded on chromosome 21 (Antonarakis, 2017; Letourneau et al., 2014; Liu et al., 2017; Stamoulis et al., 2019). It has been proposed that gene dosage compensation can ameliorate the expression levels of genes located on triplicated chromosomes, yet the mechanisms for this process remain unclear (Kojima and Cimini, 2019). Furthermore, it is thought that an extra copy of chromosome 21 has a significant impact on the rest of the genome, causing dysregulation of expression of defined chromosomal domains (Letourneau et al., 2014). To test these results, we initially focused on quantifying the mRNA expression changes of the genes on trisomic human chromosomes in primary fibroblasts. We obtained two cell lines from the Coriell Institute with trisomy 13, two with trisomy 18, seven with trisomy 21, and six euploid controls for a total of 17 cell lines (Figure S1A). All cell lines were grown in the typical tissue culture growth conditions, and samples were split into two for transcriptome and proteome quantifications. RNAseq of the transcriptomes provided quantitative information for 10,717 genes across all 17 samples (Figure 1A, Table S1). Log_2_ of the fold-changes (FC) was calculated for all 17 samples using the average expression of the six controls as the reference genome. We obtained log_2_ ratios for 92, 145, and 183 genes on chromosomes 21, 18, and 13, respectively (Figure 1B). As expected in the absence of dosage compensation, we calculated an average of approximately 1.5-fold increase (3/2 copies, log_2_ (FC) = 0.58) in gene expression of the triplicated chromosomes in all 11 trisomic cell lines analyzed (Figure 1B and S1B).

**Figure 1.**
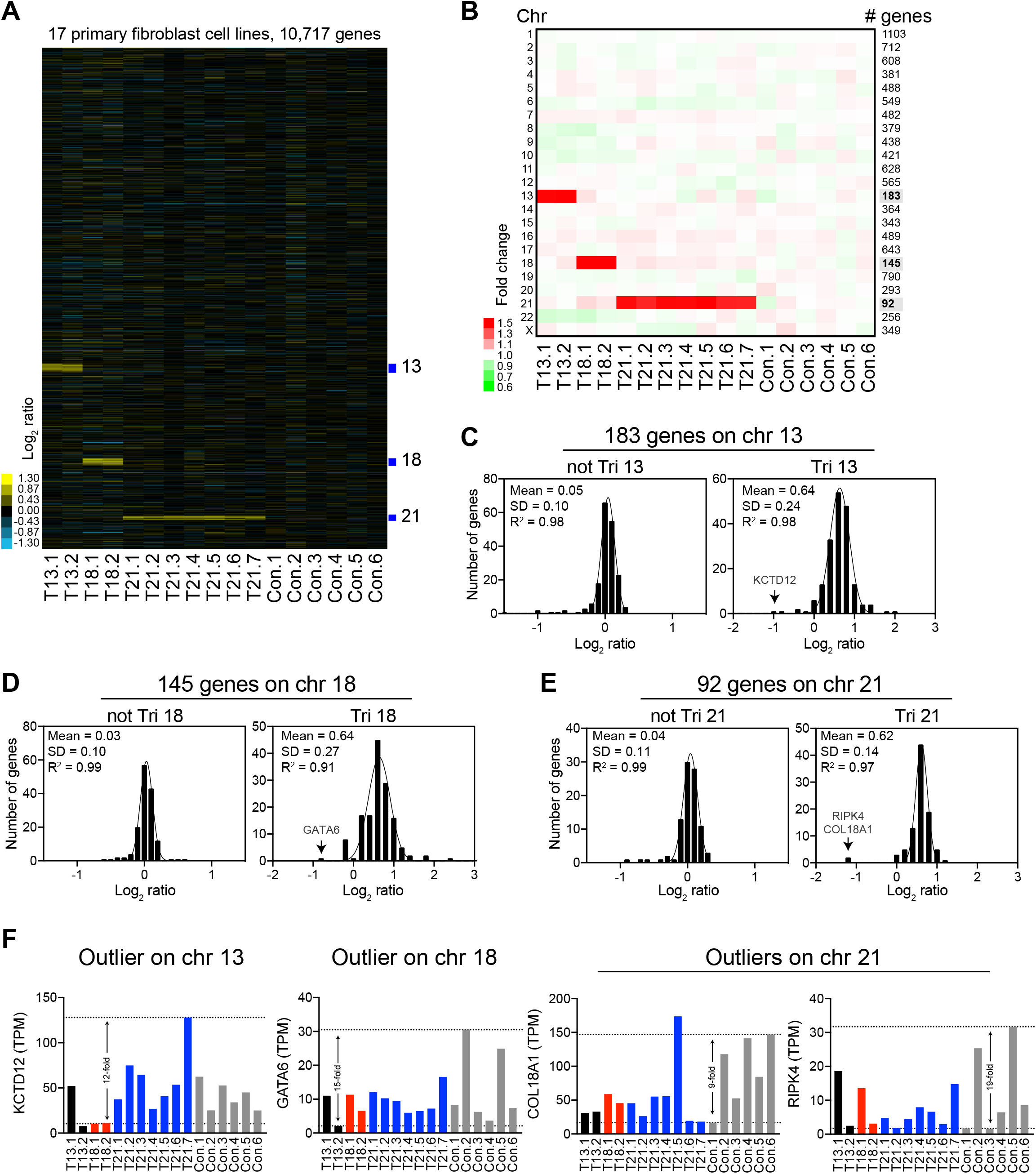
Transcript levels increase proportionally with gene copy number in trisomic primary fibroblasts. **A**. Gene expression of seventeen primary fibroblast cell lines ordered by chromosome position. Experiments (columns) for each cell line are shown. T13 = trisomy 13, T18 = trisomy 18, T21 = trisomy 21, Con = euploid control, see supplemental Figure 1A for detailed cell line nomenclature. **B**. The average gene expression per chromosome was calculated for each cell line. The number of genes quantified per chromosome is shown in the right of the heat map. **C.** Histogram of the average log_2_ ratios of the RNA copy number of genes located on euploid chromosomes (left panel) and genes present on trisomic chromosome 13 (right panel) in cell lines GM00526 and GM02948, relative to euploid controls are shown. **D.** Histogram of the average log_2_ ratios of the RNA copy number of genes located on euploid chromosomes (left panel) and genes present on trisomic chromosome 18 (right panel) in cell lines GM00734 and GM03538, relative to euploid controls are shown. **E.** Histogram of the average log_2_ ratios of the RNA copy number of genes located on euploid chromosomes (left panel) and genes present on trisomic chromosome 21 (right panel) in cell lines GM04616, GM04592, GM05397, GM06922, GM02767, GM08941, and GM08942, relative to euploid controls are shown. In C-D, the bin size for all histograms is log_2_ ratio of 0.2 and medians are identical to means. Fits to a normal distribution (black line), means, standard deviation (SD), and goodness of fit (R^2^) are shown for each distribution. **F.** Examples of the normalized transcript count (TPM, transcripts per million) of a couple of triplicated genes that are outliers in the fits of the normal distributions. Black, red, blue, and gray bars correspond to expression in trisomy 13, 18, 21, and controls, respectively.

The values of the average log_2_ ratios of the triplicated genes fit a normal distribution with means of 0.6 and standard deviations (SD) of 0.24, 0.27, and 0.14 for trisomies 13, 18, and 21, respectively (Figures 1C-E). Although the log_2_ ratios fit a normal distribution quite well (R^2^s between 0.91-0.98), there are a couple of gene outliers whose expression does not appear to increase with copy number. Closer inspection of the normalized transcript counts (TPM, Transcript Per Million) of the outliers in each distribution revealed that their expression levels show high variability across individual cell lines, including the six control cell lines (Figure 1F and Figure S1C). For example, one outlier encoded on chromosome 13 is KCTD12, a gene whose expression varies up to 12-fold among cell lines and up to 7-fold in the two different trisomies 13. The levels of the transcription factor GATA6 encoded on chromosome 18 vary up to 15-fold among cell lines not trisomic for chromosome 18. COL18A1 and RIPK4 are the main outliers in trisomy 21 and show 9-fold and 19-fold, respectively, variability among control cell lines. Indeed, most outliers of the distributions of triplicated genes, including below or above the fits, show high variability in expression among different individuals (Figure S1C). Our analysis indicates that, on average, most genes located on the trisomic chromosomes increase 1.5-fold in expression and that interindividual variability in gene expression accounts for a couple of outliers in the dataset.

Several recent studies of the transcriptome profiles of trisomy 21 cells have been performed, but a precise quantification and analyses of the mRNA levels of genes encoded on chromosome 21 were not reported. To validate that transcript levels of chromosome 21 genes proportionally increase with copy number, we analyzed two recently published datasets, one by Sullivan et al. (Sullivan et al., 2016) and the other by Letourneau et al. (Letourneau et al., 2014). In a tour de force approach, Sullivan et al. obtained transcriptome profiles of 24 hTERT immortalized fibroblasts, 12 transformed B cells (lymphoblastoid cell lines, LCLs), 17 monocytes, and 17 T cells lines from different individuals with normal karyotype or with Down syndrome (Figure S2A). We calculated the log_2_ of the FC using the average expression of the control individuals as the reference genome for each cell type and plotted the data ordered by chromosome position (Figure 2A, Table S2). Consistent with our analysis of primary fibroblasts, we found that on average, the mRNA levels of genes located on chromosome 21 show a 1.5-fold increase in all cell types and across samples from different individuals with Down syndrome relative to the controls (Figure 2A-B). As observed in our data, a few outliers can be accounted for by their high variability in expression levels among control individuals (Figure S2C). Intriguingly, one fibroblast cell line (T21.1b) shows significant changes in gene expression in all chromosomes and deviates from the rest. We suspect that technical issues may account for such extreme changes in gene expression across the genome in this sample (see below). Altogether, the expression of chromosome 21 genes in 37 out of 38 cell lines with trisomy 21 analyzed by Sullivan et al. fit a normal distribution with means very close to the predicted log_2_ ratio of 0.58 (Figure 2B).

**Figure 2.**
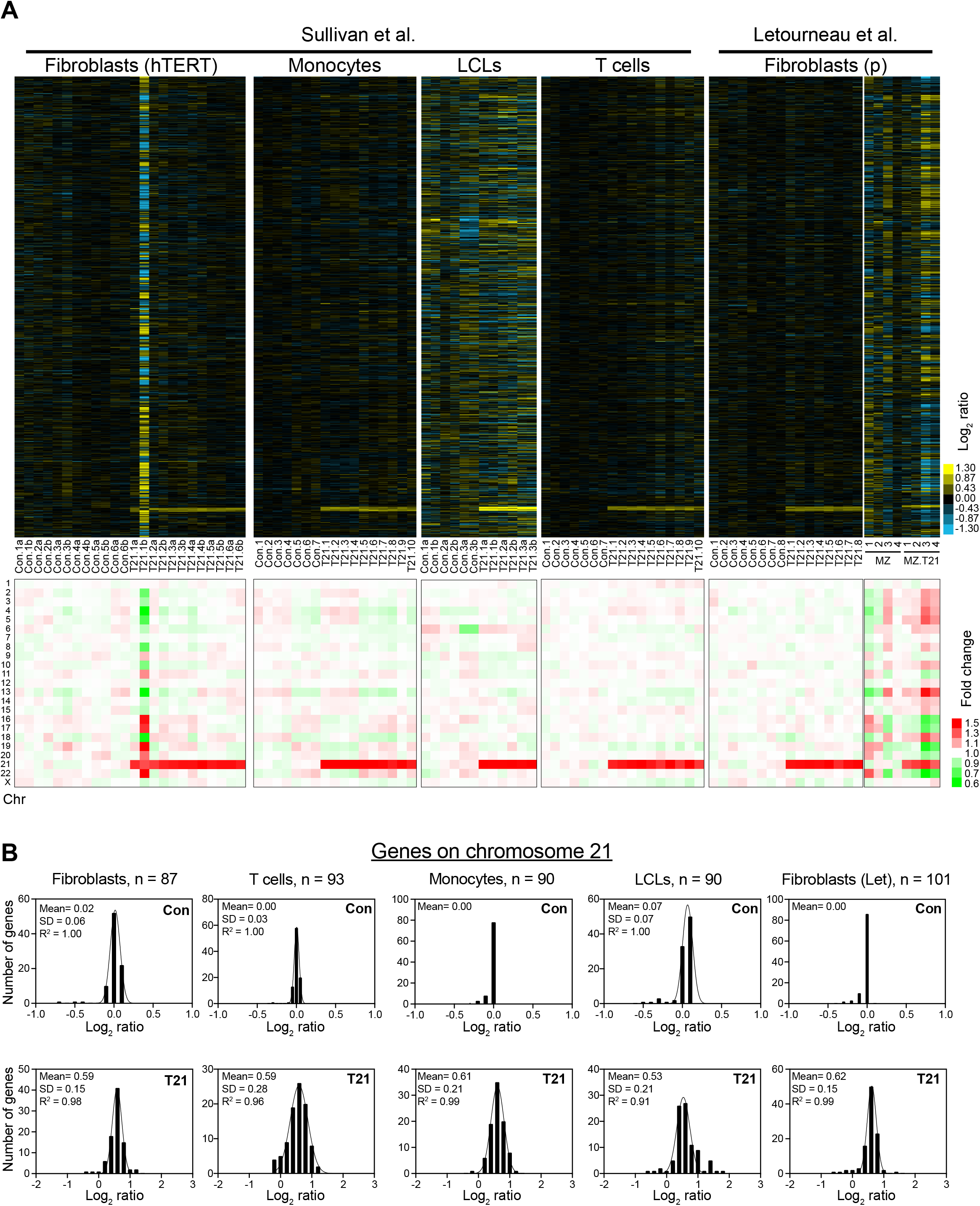
Transcript levels increase proportionally with gene copy number in distinct trisomy 21 cell lines. **A**. Gene expression of 24 immortalized fibroblast cell lines, 17 monocytes cell lines, 12 lymphoblastoid cell lines (LCLs), 17 T cell lines obtained by Sullivan et al., and 18 primary fibroblast and 4 technical replicates of primary fibroblasts from monozygotic twins obtained by Letourneau et al. Experiments (columns) for each cell line are shown ordered by chromosome position. See supplemental Figure 2A for detailed cell line nomenclature. Bottom, average gene expression per chromosome was calculated for each cell line. **B.** Histogram of the average log_2_ ratios of the RNA copy number of genes located on euploid chromosomes (top panels) and genes present on trisomic chromosomes (top panels), of the trisomic cell lines analyzed in **(A)** relative to euploid controls are shown. Bin size for all histograms is log_2_ ratio of 0.2 and medians are identical to means. Fits to a normal distribution (black line), means, standard deviation (SD), and goodness of fit (R^2^) are shown for each distribution.

In the second study, Letourneau et al. reported the expression profiles of primary fibroblasts from 16 unrelated individuals (8 controls and 8 with trisomy 21), and of two cell lines obtained from monozygotic twins, one with a normal karyotype and the other with trisomy 21 (Figure S2B). Consistently, we found that the expression of the primary fibroblasts from individuals with Down syndrome shows, on average, a 1.5-fold increase on chromosome 21. On the other hand, the monozygotic twin with trisomy 21 shows very variable expression across the genome (Figure 2A, Table S2). Letourneau et al. interpreted the latter result as evidence for trisomy 21 driving the dysregulation in expression across chromosomal domains. A closer inspection of the transcript counts raises the possibility that technical issues account for this result. We found that the correlation of gene expression among biological replicates of the same cell line from the monozygotic twin with trisomy 21 is poor (R^2^ = 0.44) compared to the correlation between primary fibroblasts from different individuals (R^2^ = 0.95, Figures S2D-E). We also found that up to 20% of the total reads in the RNAseq data of the monozygotic twin come from mitochondrial DNA encoded genes (Figure S2E). Unexpectedly, the average gene expression per chromosome of the outlier sample from the Sullivan et al. study (T21.b) shows similar expression patterns but with different values compared to the monozygotic twin with trisomy 21 (Figure S2F). This analysis raises the possibility that similar technical issues may account for these very variable transcriptomes. Excluding the two outlier transcriptomes, our analyses support the hypothesis that, on average, the mRNA levels increase proportionally with gene copy numbers. Importantly, mechanisms to compensate and ameliorate gene expression of an extra copy of a human autosome do not seem to be engaged.

### Interindividual variability drives gene expression patterns of human primary fibroblasts

Similar to the findings reported by Sullivan et al. using principal component analysis, we found that the gene expression patterns of human fibroblasts do not cluster with karyotype, sex, or the donor’s age in all three datasets (Figure 3A). Excluding chromosome 21, we were not able to identify a set of genes that are commonly up or downregulated in all three datasets. Instead, hierarchical clustering analysis indicates that several control cell lines show gene expression patterns closer to a trisomic cell line than to another control. These results suggest that gene expression patterns are specific to the identity of the cell coming from different individuals causing significant variability, thereby occluding the identity of specific gene responses to aneuploidy in this context.

**Figure 3.**
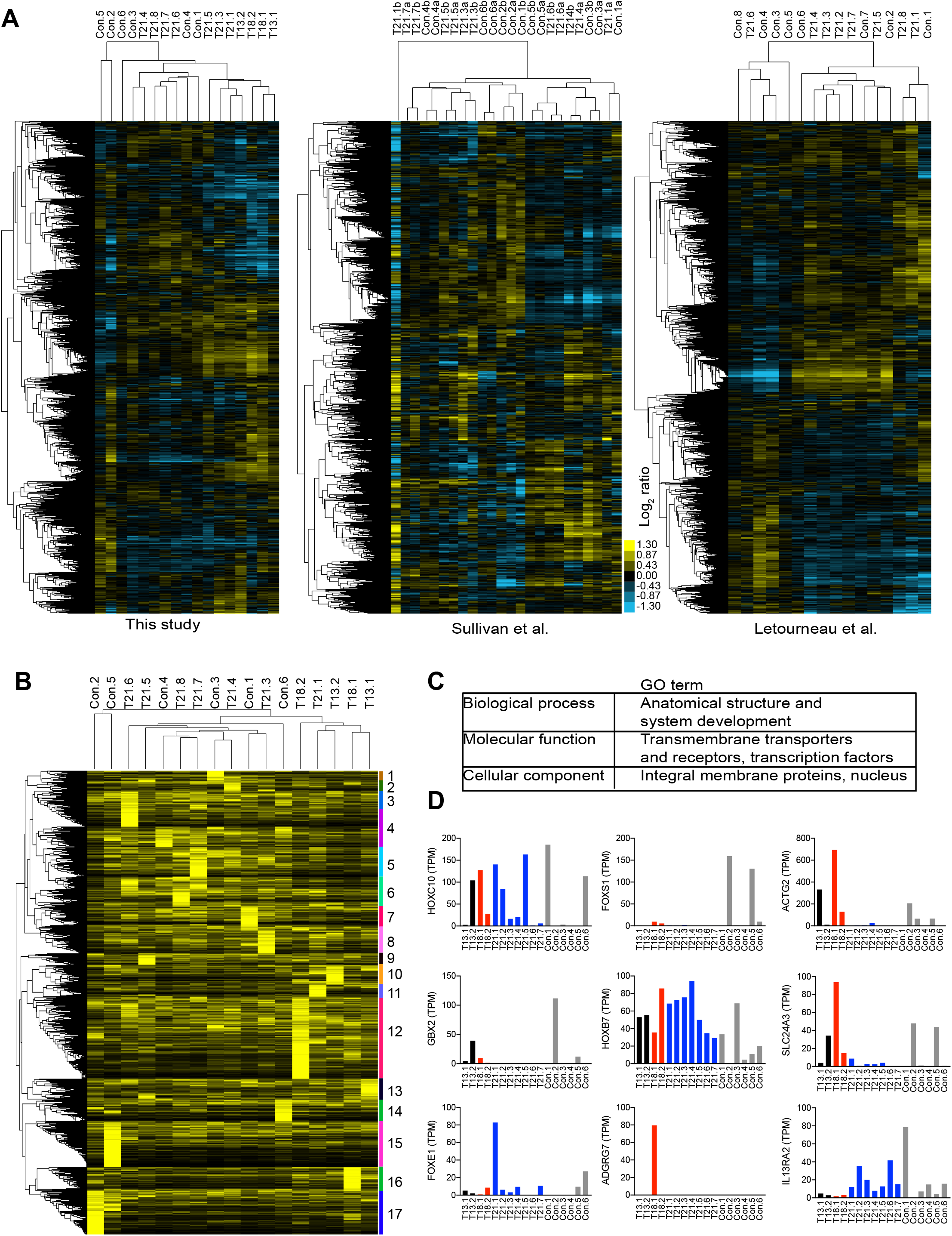
Interindividual variability drives gene expression patterns of human primary fibroblasts. **A.** Hierarchical clustering analysis of the expression patterns of primary fibroblasts analyzed in this study, in the studies by Sullivan et al. and the Letourneau et al. are shown. Expression patterns cluster independent of the karyotype of the cell line. **B.** Relative gene expression of genes uniquely expressed in some cell lines. Hierarchical clustering analysis revealed 17 clusters specific for each cell line. **C.** Gene ontology enrichment analysis shows that the uniquely expressed genes are enriched in master transcription factors that regulate development and cell surface markers, including transmembrane transporters and receptors proteins. **D.** Examples of the normalized transcript count (TPM, transcripts per million) of a couple of genes highly expressed in some cell lines but poorly expressed in others. Black, red, blue, and gray bars correspond to expression in trisomy 13, 18, 21, and controls, respectively.

To better define the source of variability in gene expression among primary fibroblasts, we analyzed the identity of genes uniquely expressed in individual cell lines. Initially, genes whose expression was not detected (zero reads) in several cell lines were omitted as part of our cutoff to calculate FC among all samples. We found a total of 2,543 that were significantly expressed in specific cell lines but not detected in others (Table S3). Rather than calculating ratios, we normalized the relative T.P.M. across genes from 0 reads to maximum reads to a scale range from 0 to 1 (0 is in black, 1 is in yellow, Figure 3B). Hierarchical clustering revealed a set of genes specifically expressed in each cell line. We identified 17 clusters representing 17 cell lines. Remarkably, the clustering pattern among the uniquely expressed genes in the cell lines is similar to the clustering pattern of the gene expression of their whole transcriptome (Figures 3A-B). Gene ontology enrichment analyses revealed that this set of genes includes several master transcription factors that determine the anatomical structure and regulate development, including several homeobox (HOX) and forkhead box (FOX) genes (Figure 3C-D). Several membrane transporters and receptors are present in this set of genes. These results indicate that cell identity, including anatomical and developmental properties differing in each individual, governs the gene expression patterns of primary fibroblasts in both trisomic and euploid cell lines.

Other sources of variability on gene expression come from culture conditions. We found that biological replicates of the same cell line cultured in parallel show good reproducibility (linear regression fit, slope = 1.0 and R^2^ = 0.97, Figure S3B). However, when we compared the gene expression of the same cell lines grown months apart using independent media preparations, hierarchical clustering patterns were influenced by the time the cultures were performed (Figure S3C-D). These results indicate that in addition to cell identity, specific growth conditions can significantly contribute to physiological noise in gene expression, which may fuel the discrepancies in the results between different research groups.

Despite the interindividual variability, we sought to determine whether there is a set of genes that change expression in response to aneuploidy or trisomy 21 in our dataset. Using hierarchical clustering analyses, we identified a set of commonly down-regulated genes in 7 out of 11 trisomic cell lines (Figure S3A). This cluster is enriched for genes that regulate metabolic processes, including amino acid metabolic process (p-value = 6.8 E-6) and response to nutrient levels (p-value = 7.1 E-7). These results support the hypothesis that aneuploidy disrupts cellular metabolism. Next, by sorting the average change in gene expression between aneuploids compared to euploids, we also identify a set of upregulated genes in trisomic fibroblasts. The upregulated cluster is enriched for genes in the DNA repair (p-value 6.2 E-21) pathway, consistent with aneuploidy causing replication stress and genomic instability. Noteworthy, Sullivan et al. found that trisomy 21 fibroblasts show a gene expression pattern associated with an activated immune response driven by interferon signaling, a result not observed in the analysis of our dataset. Furthering these discrepancies, Letourneau et al. reported another signature associated with reduced expression of secreted proteins involved in cytokine-cytokine receptor pathways and inflammatory response (Letourneau et al., 2014). Altogether, the three datasets revealed a different set of genes that might be regulated by trisomy 21 without reaching a consensus on a conserved cellular response. Other studies have also reported different cellular pathways affected by trisomy 21 (Gonzales et al., 2018; Scarpato et al., 2014; Zamponi et al., 2018). We conclude that interindividual variability and experimental conditions mask gene expression patterns that might be associated with cellular responses associated with aneuploidy.

### Protein levels increase proportionally with gene copy number and mRNA levels

Next, we analyzed how changes in mRNA levels modify the proteome. We used isobaric tandem mass tag (TMT)-based quantitative mass spectrometry to quantify protein levels in primary fibroblasts. We analyzed two sets of 10 samples using a TMT 10-plex protocol. One control (Con.1) was analyzed in duplicate in both sets of samples to monitor our approach’s technical variability. Also, samples for one trisomy 21 cell line (T21.1) were included in both sets of quantifications to test reproducibility. We obtained quantitative information for 7,273 proteins and 7,297 proteins in the first and second sets of analysis, respectively, with 6,486 proteins overlap (Figure 4A, Table S4). A total of five controls, two trisomies 13, two trisomies 18, and seven trisomies 21, proteomes were obtained in our analysis. A comparison of the peptide counts of the two control samples (Con.1a and Con.1b) shows good reproducibility with a linear correlation of slope equal to 1 (R^2^ = 0.99); while the comparison of the proteomes of two unrelated individuals shows more variability (R^2^ = 0.81, Figure 4B).

**Figure 4.**
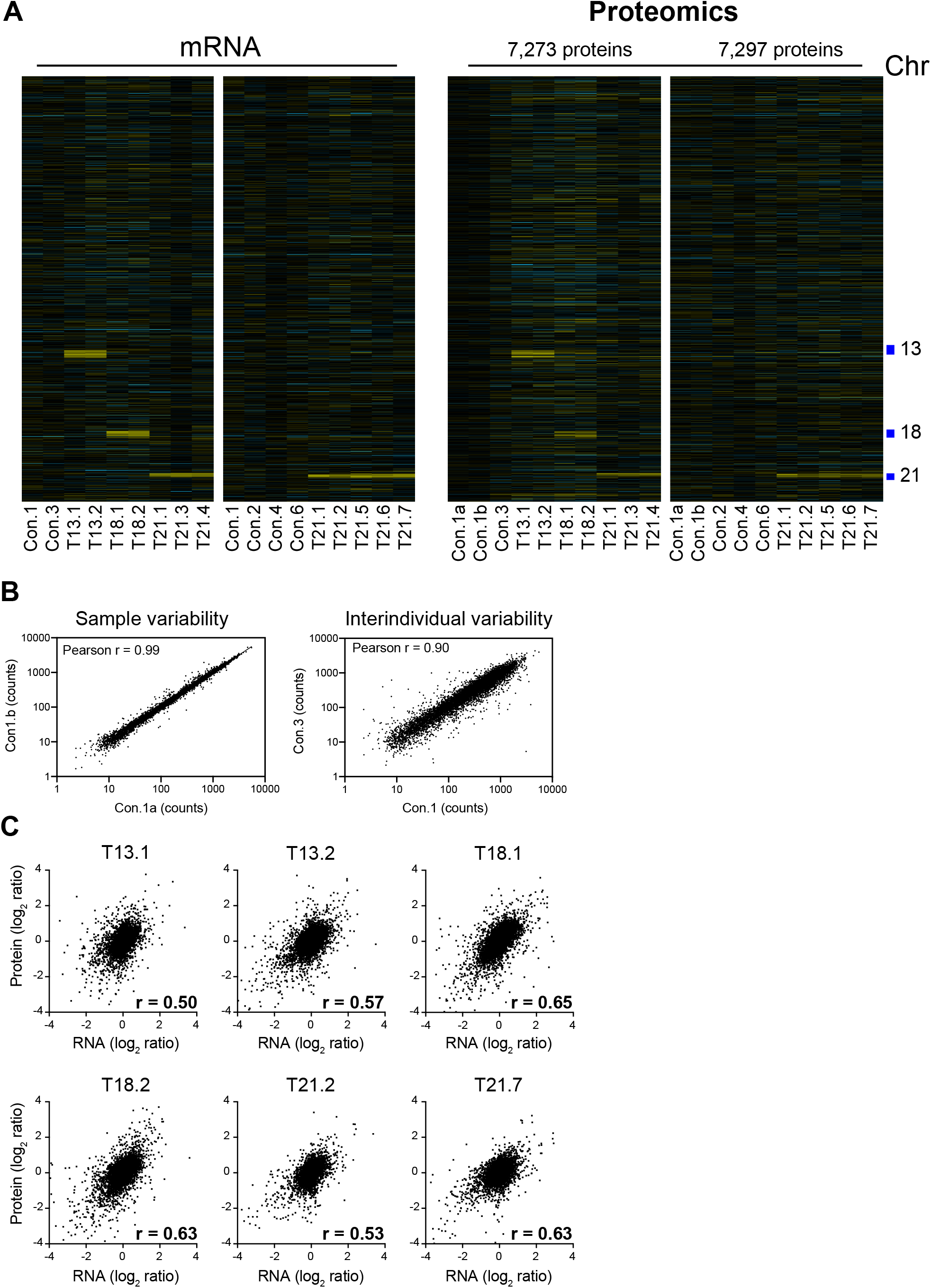
Protein levels proportionally increase with copy number in trisomic fibroblasts. **A.** Comparison of the mRNA (left) and protein levels (right) of human primary fibroblasts. Genes are ordered by chromosome position in each experiment (columns). **B.** Linear regression analysis of protein counts in technical replicates of a control cell line and between two cell lines from different individuals. **C.** Pearson correlation r was calculated for the mRNA and protein levels for six representative trisomic cell lines.

We then calculated the log_2_ of the FC using the average counts of the control samples as the reference proteome in each dataset. Increases in protein levels readily reveal the identity of the triplicated chromosomes by plotting of the proteome profiles for each cell line ordered by chromosome position (Figure 4A). Importantly, changes in RNA and protein levels correlate with coefficients (Pearson r) ranging from 0.4 to 0.7 among all samples analyzed. Indeed, proteome profiles could easily be matched to their corresponding RNA profiles using correlation analysis between log_2_ ratios of RNA and protein in all samples (Figure S4A). 129, 110, and 73 proteins on chromosomes 13, 18, and 21, respectively, were quantified in the first dataset, while 76 proteins on chromosome 21 were quantified in the second set. On average, the protein levels of the triplicated chromosomes increase by log_2_ ratios of 0.45, 0.43, and 0.40 in trisomy 13, 18, and 21, respectively (SD between 0.3-0.5, Figure 5A and 5B). These results indicate the protein levels proportionally increase with copy number but are slightly lower (1.3 to 1.4-fold) than the theoretical 1.5-fold increase observed in transcript levels. The distributions of the log_2_ ratios fit quite well a normal distribution (R^2^s between 0.90-0.98) with a couple of outliers both below and above the curve. As observed in the transcript levels, closer inspection of the outliers indicates that interindividual variability accounts for their apparent dysregulation (a few examples are shown in Figure S4B).

**Figure 5.**
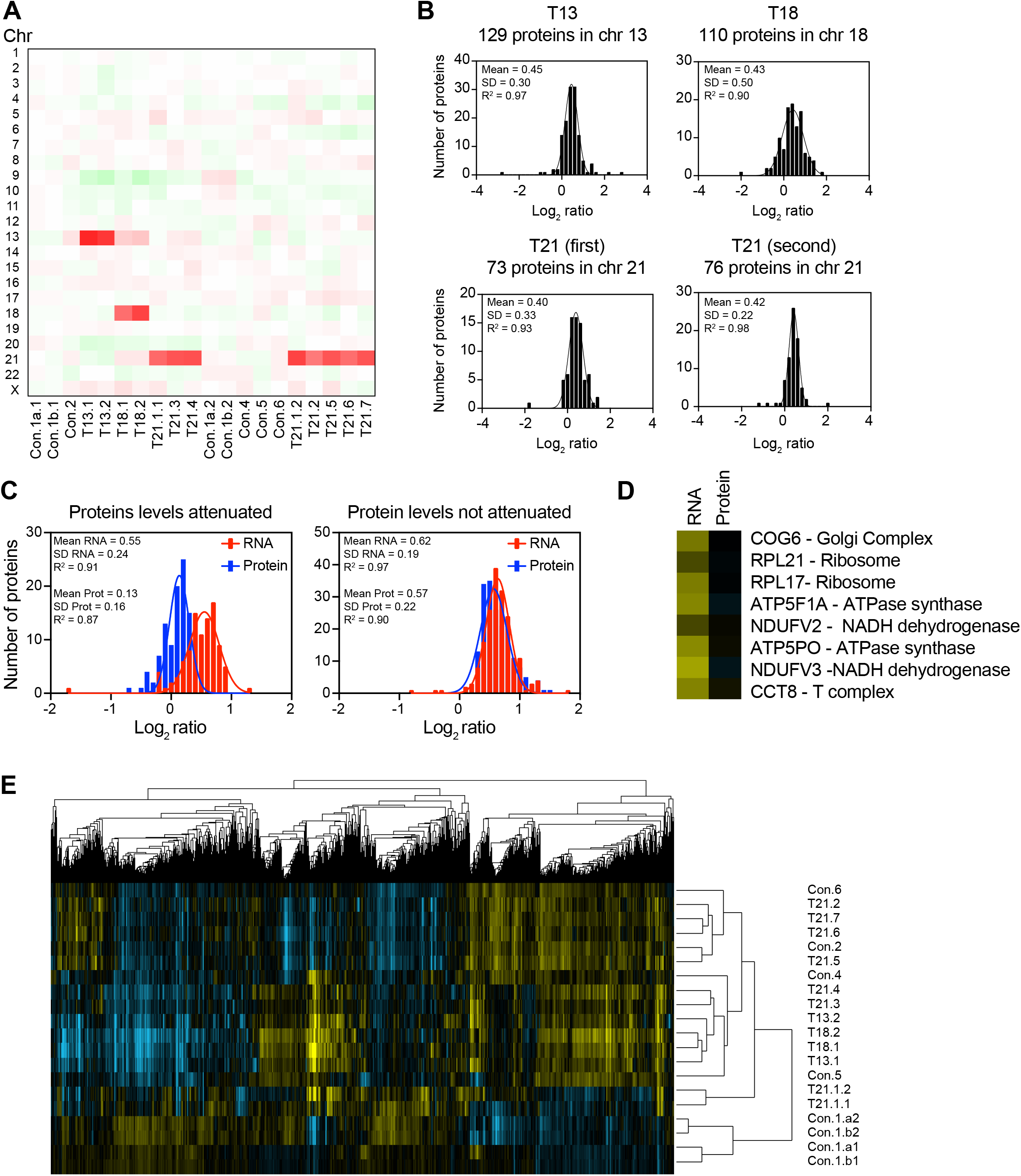
Protein levels of subunits of macromolecular complexes are attenuated in trisomic primary fibroblasts. **A.** Heatmap of the average protein levels per chromosome in primary fibroblasts. Experiments (columns) for each cell line are ordered by chromosome position. **B**. Histogram of the average log_2_ ratios of the protein levels of genes located on chromosome 13 in trisomy 13 cell lines (top left), on chromosome 18 in trisomy 18 cell lines (top right), on chromosome 21 in trisomy 21 cell lines in the first (lower left) and second (lower right) datasets, relative to euploid controls are shown. The bin size for all histograms is log_2_ ratio of 0.2. Fits to a normal distribution (black line), means, standard deviation (SD), and goodness of fit (R^2^) are shown for each distribution. **C.** Histogram of the average log_2_ ratios of the RNA (red) and protein (blue) levels of triplicated genes that show protein levels lower than predicted (left) and 1.5-fold changes in trisomic fibroblasts. The bin size for all histograms is log_2_ ratio of 0.1. Fits to a normal distribution (solid lines), means, standard deviation (SD), and goodness of fit (R^2^) are shown for each distribution **D.** A few examples of individual subunits located on triplicated chromosomes that show attenuation at the protein levels compared to RNA. COG6 and RPL21 are in chromosome 13. RPL17, ATP5F1A, and NDUFV2 are in chromosome18. ATP5PO, NDUFV3, and CCT8 are in chromosome 21. **E.** Hierarchical clustering analyses of proteome profiles do not cluster by karyotype of the cells lines.

### Subunits of macromolecular complexes are regulated post-transcriptionally

To better understand why the fold increase of the average protein levels of triplicated chromosomes is slightly lower than the predicted 1.5-fold, we analyzed the identity of a set of proteins encoded on the triplicated chromosomes with lower levels than expected in the trisomic cell lines. Excluding the outliers due to interindividual variability, we found that about one-third of triplicated genes (97 out of 281) show protein changes of log_2_ ratios less than 0.3 and fit a normal distribution with a mean of 0.13 (SD = 0.16, Figure 5C). Remarkably, the average change in mRNA levels of this set of proteins is 1.5-fold (mean log_2_ ratio = 0.55, Figure 5C). These results indicate that the levels of this set of proteins are attenuated via posttranscriptional mechanisms. The other 184 triplicated genes show the predicted average increase of 1.5-fold in both mRNA and protein levels (Figure 5C).

We and others showed that a set of attenuated proteins in aneuploid yeast and mammalian cell lines include subunits of macromolecular complexes (Dephoure et al., 2014; McShane et al., 2016; Stingele et al., 2012; Torres et al., 2007). We hypothesized that protein instability of individual subunits in the absence of the upregulation of entire complexes is the cause of this effect (Dephoure et al., 2014). Here, we found that 39% (38 of 97) of attenuated proteins are subunits of macromolecular complexes. Notably, 2 out of 2 ribosomal subunits in our dataset show attenuation consistent with the finding that ribosomal subunits are rapidly degraded unless they assemble into a stable complex (Dephoure et al., 2014; elBaradi et al., 1986; Tsay et al., 1988). Other examples include 2 subunits of the ATPase synthase and 2 subunits of NADH dehydrogenase (Figure 5D). We also found that several membrane proteins and secreted proteins account for the attenuation in protein levels. Consistent with previous studies in aneuploid yeast cells, our results indicate that protein levels increase in abundance proportionally with mRNA levels with a set of proteins being attenuated by posttranscriptional mechanisms most likely mediated by protein degradation pathways. Therefore, a conserved consequence of aneuploidy in trisomy 21 cells is increased protein synthesis leading to an imbalanced proteome and higher activity of the protein quality control pathways.

### Interindividual variability determines proteome profiles of human primary fibroblasts

Hierarchical clustering analysis also shows that the proteome profiles cluster independent of karyotype (Figure 5E). Several proteome profiles of controls are more similar to trisomic proteomes than to other controls. For example, the proteome profile of Con.2 is more similar to T21.6, T21.7, and T21.2 than to the proteome of Con.1. Importantly, we found a significant correlation between the proteome profiles of the replicates within each dataset (Con.1a vs. Con.1b) and in between datasets (T21.1.1 vs. T21.1.2) indicating good reproducibility of our analysis.

Hierarchical clustering did not reveal a clear pattern of commonly up- or down-regulated proteins in the aneuploid cell lines. Since there is a strong correlation between RNA and protein changes, we analyzed the protein levels of the down and upregulated gene clusters identified in the transcriptome analyses (Figure S5). Consistently, we found that these clusters show specific down and upregulation at the protein levels too. However, analysis of the average changes shows that these responses are significantly attenuated. Importantly, this set of proteins remains enriched for factors that regulate amino acid biosynthesis in the downregulated cluster and DNA replication in the upregulated cluster.

### Trisomic fibroblasts rely on serine-driven lipid synthesis to proliferate

Altered sphingolipid metabolism is a conserved aneuploid-associated phenotype of aneuploid yeast and mammalian cells (Hwang et al., 2017; Tang et al., 2017). Sphingolipids are essential molecules that are integral structural components of cellular membranes. We recently showed that altered sphingolipid levels are associated with abnormalities in the morphology of the nucleus of aneuploid cells, including human primary fibroblasts trisomic for chromosomes 13, 18, or 21 (Hwang et al., 2019). Furthermore, we found that sphingolipid levels in human trisomic fibroblasts are elevated compared to euploid cells. The pathway of the synthesis of sphingolipids is conserved in yeast and humans. The initial step involves the condensation of the amino acid serine to palmitoyl-CoA by serine palmitoyltransferase (SPT) to generate long-chain bases (Figure 6A). Analysis of the mRNA levels of trisomic cell lines revealed that the expression of one of the central subunits of SPT, SPTLC2, is upregulated in several trisomic fibroblasts cell lines compared to controls in our data set (Figure 6B). Notably, the upregulation of the SPTLC2 mRNA is also detected in several trisomy 21 cell lines analyzed in the Sullivan et al. and Letourneau et al. studies (Figure 6B-C). The SPTLC2 upregulation could lead to increased SPT activity as this subunit is required for the stability of the enzyme (Yasuda et al., 2003). Consistently, our proteome analyses indicate that the levels of SPTLC2 increase in trisomic fibroblasts compared to controls (Figure 6D). To validate these results, we performed western blot analysis of SPTLC1 and SPTLC2 in two pairs of controls, trisomy 13, 18, and 21, and found that SPT’s protein levels are increased by aneuploidy. These results provide a mechanism by which the synthesis of sphingolipids is upregulated in aneuploid cells. Further, they suggest that the regulation of the mRNA levels one of the main subunits can stabilize the complex and increase enzyme levels in the cells.

**Figure 6.**
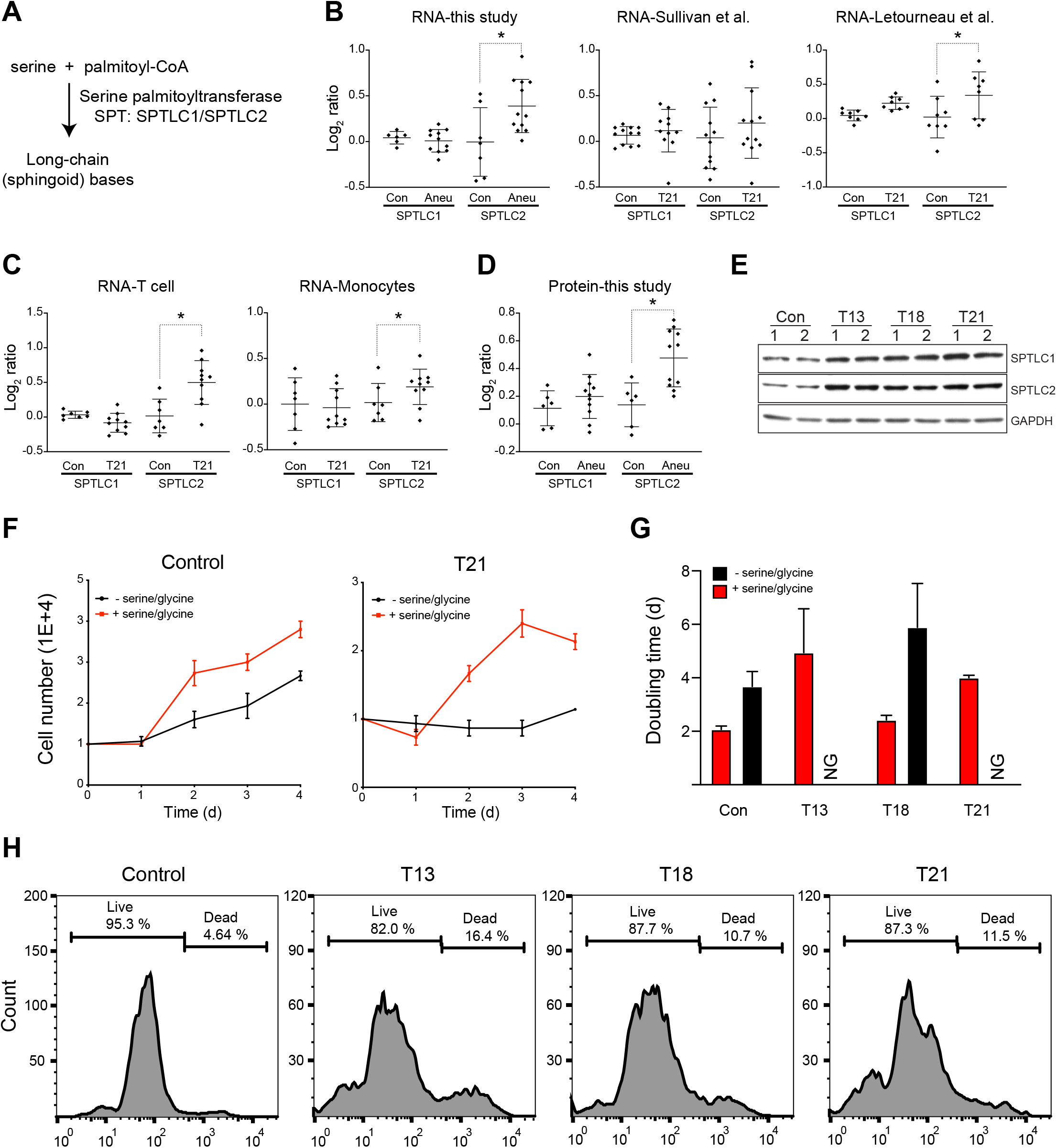
Trisomic primary fibroblasts rely on serine-driven lipid biosynthesis and show lower viability compared to euploid controls. **A.** Schematic of the biochemical pathway for the serine-dependent synthesis of sphingolipids. **B.** Expression levels of subunits SPTLC1 and SPTLC2 of the serine palmitoyltransferase enzyme in fibroblasts in this study, and in the Sullivan et al. and Letourneau et al. studies. **C.** Expression levels of subunits SPTLC1 and SPTLC2 in T cells and Monocytes. **D.** Protein levels of SPTLC1 and SPTLC2 in primary fibroblasts. **E.** Western blots of SPTLC1 and SPTLC2 in primary fibroblasts. GADPH was used a loading control. **F.** Proliferation of control fibroblasts (left) and trisomy 21 fibroblasts (right) with or without serine and glycine. Glycine was depleted because it can be used to generate serine by the SHMT1/2 enzymes**. G.** Doubling times of primary fibroblasts with or without serine/glycine. **H.** Viability assay shows that there is an increase in cell death in the trisomic fibroblasts.

Sphingolipid synthesis is tightly coupled to the availability of intracellular serine, and aneuploid yeast cells strictly rely on serine synthesis to proliferate (Cowart and Hannun, 2007; Hwang et al., 2017). Here, we tested whether trisomy fibroblasts rely on serine to proliferate. Remarkably, we found that trisomy 13 and 21 fibroblasts are not able to proliferate in media depleted of serine while the proliferation of trisomy 18 was significantly more affected compared to controls (Figure 6F-G). These results indicate that serine, an amino acid utilized for protein, lipid, and nucleotide biosynthesis, is essential for the proliferation of trisomy 21 cells. These results support the hypothesis that aneuploidy disrupts cellular metabolisms due to a higher metabolic demand driven by increased nucleotide, protein, and lipid synthesis.

### Aneuploidy lowers the viability of human primary fibroblasts

Previous studies have shown that mammalian trisomic cells proliferate slower than their euploid counterparts (Hwang et al., 2019; Stingele et al., 2012; Torres et al., 2010; Torres et al., 2007; Williams et al., 2008), an observation reported almost 50 years ago for trisomy 21 cells (Segal and McCoy, 1974). However, the mechanisms behind the slow proliferation phenotype remain unknown. Early passage trisomic cells show no signs of cell cycle arrest or senescence assayed by beta-galactosidase staining. As reported in trisomic mouse embryonic fibroblasts (MEF) (Williams et al., 2008), we could not detect significant cell cycle delays in human trisomic fibroblasts. Since cell cycle delays or senescence do not account for lower proliferation rates of trisomic cells, and because aneuploidy lowers the viability of yeast cells (Torres et al., 2007), we posit that cell viability must be affected in trisomic fibroblasts. To investigate whether cellular viability is affected by aneuploidy, we used the LIVE/DEAD™ Fixable Green Dead Cell Stain Kit. This kit uses a dye that reacts with free amines yielding intense fluorescent staining. In viable cells, the dye’s reactivity is restricted to the cell surface resulting in less intense fluorescence than non-viable cells wherein the dye diffuses inside the cell (Figure S6). Remarkably, we found that human trisomic fibroblasts show a significant increase in the population of non-viable cells. While control cell lines show up 4%, trisomy 13, 18, or 21 show 16%, 11%, and 12% of non-viable cells, respectively (Figure 6H). Such decreases in viability can translate into a significant difference in cell numbers during exponential growth. Our results indicate that lower viability is a major consequence of trisomy 21 due to the cell’s aneuploid status and that this contributed to the low proliferation rates observed in vitro.

## Discussion

The acquisition of an extra autosome has detrimental consequences to organismal development. At the cellular level, studies in yeast indicate that the deleterious effects of aneuploidy increase with the size of the extra chromosome (Torres et al., 2007; Torres et al., 2008). In humans, three copies of chromosomes 21, 18, or 13, which encode for the least number of genes in that order, are the only viable trisomies. Notably, the effects on human development in trisomies 13 and 18 are more deleterious than trisomy 21. This fact is consistent with a correlation between the number of genes on the triplicated chromosome and the degree by which aneuploidy disrupts cellular physiology. In the context of Down syndrome, a genecentric view poses that increased expression of the triplicated genes encoded on chromosome 21 causes cellular defects (Antonarakis et al., 2004; Antonarakis et al., 2020). Our studies show that trisomy 21 cells shared several phenotypes with other mammalian aneuploid cells, including trisomic MEFs, human trisomies 13 and 18, and even with aneuploid yeast cells (Hwang et al., 2019; Pfau and Amon, 2012; Stingele et al., 2012; Torres et al., 2007; Williams et al., 2008). These aneuploidy-associated phenotypes across organisms and independent of the triplicated chromosome identity include hampered proliferation, genomic instability, abnormal nuclear morphology, and altered metabolism. Therefore, to better understand the pathophysiology of individuals with Down syndrome, aneuploidy’s effects on the physiology of the cell must be taken into account.

Trisomies associated with sex chromosomes, such as the triple X and XYY syndromes, are not as deleterious to human development as autosomal trisomies. In the case of triple X, the non-coding RNA Xist silences the expression of the third copy of the X chromosomes. Since the X chromosome is larger in sequence than chromosomes 21, 18, or 13, the mere presence and maintenance of extra DNA are not significant drivers of aneuploidy-associated phenotypes. In support of this, yeast harboring yeast artificial chromosomes with heterochromatic human DNA comparable in size to the yeast chromosomes do not show aneuploidy-associated phenotypes (Hwang et al., 2017; Torres et al., 2007). In the case of an extra copy of the Y chromosome, the number of genes expressed and encoded on the Y chromosome is significantly smaller than that of chromosome 21. Together, these observations support the hypothesis that increases in gene expression must be the cause of cellular imbalance and the disruption of cellular homeostasis.

Several studies have suggested that dosage compensation could ameliorate the RNA levels of the genes present on an extra copy of an autosome (Disteche, 2016; Kojima and Cimini, 2019). However, mechanisms for dosage compensation of an entire extra copy of an autosome are not known in human cells. Here, we show that on average, mRNA levels increase 1.5-fold in human cells trisomic for chromosomes 13, 18, or 21, and our analysis supports the lack of dosage compensation of human autosomes. Furthermore, quantitative proteomics indicates that increases in transcript levels lead to proportional increases in protein levels except for a subset of proteins that encode subunits of macromolecular complexes. Supported by several studies, individual subunits of macromolecular complexes are unstable unless they are assembled into a stable complex (Brennan et al., 2019; Dephoure et al., 2014; McShane et al., 2016; Stingele et al., 2012). Excess subunits are either degraded or aggregated, as protein aggregation is as effective as protein degradation at lowering levels of excess proteins (Brennan et al., 2019). Importantly, evolutionary mechanisms are in place to synthesize and assemble subunits of a particular complex in equimolar amounts, presumably sparing the cell from the wasteful synthesis of individual subunits (Li et al., 2014; Shiber et al., 2018). Secreted and membrane proteins are other classes of proteins that show lower levels than the predicted 1.5-fold. The secreted proteins will not accumulate in the cell but can lead to physiological changes by modifying the extracellular environment. Membrane protein levels may depend on mechanisms that control protein trafficking and endocytosis. Increased protein synthesis, folding, and degradation may affect protein quality control pathways and altered the metabolic demands of trisomy 21 cells, independent of the identity and function of the triplicated genes.

Analysis of different cell types from different individuals with and without trisomy 21 indicates that interindividual variability is a major driver of gene expression patterns. Transcriptome profiles cluster independent of karyotype, sex, or age of the donor. Our analyses of the transcriptomes of primary fibroblasts from several studies indicate that each cell line uniquely expresses a set of transcription factors associated with anatomical structure and development. To circumvent this issue, Letourneau et al. analyzed the expression pattern of two fibroblast cell lines isolated from monozygotic twins, one of them trisomy for chromosome 21. Those studies concluded that trisomy 21 leads to genome-wide disruption of chromosomal domains or territories. Two previous publications have questioned the findings by Letourneau et al. (Ahlfors et al., 2019; Do et al., 2015). Our analysis indicates that the gene expression profiles on the monozygotic cell lines are noisy, and we suspect that this is due to technical issues. Lastly, gene expression analyses of trisomic MEFs and human aneuploid cell lines generated by different approaches do not converge to a common gene expression pattern in response to aneuploidy (Santaguida et al., 2017; Stingele et al., 2012; Williams et al., 2008). In isogenic aneuploid yeast, we showed that changes in gene expression are associated with slow proliferation (Torres et al., 2007). The fact that cell cycle delays are not present in primary human cells is consistent with the lack of a transcriptional response associated with slow proliferation.

Other aneuploidy-associated phenotypes in human trisomy cells include increased replication stress measured by foci positive for the p53 Binding Protein 1(Hwang et al., 2019; Passerini et al., 2016). Mutations in genes involved in DNA replication and repair cause premature aging (Kubben and Misteli, 2017). Thus, aneuploidy driven genomic instability may play an active role in the premature aging characteristics of individuals with Down syndrome. Importantly, trisomic MEFs and human fibroblasts do not show significant defects in cell cycle progression or increased senescence. Here we found that reduced proliferation measured in cell accumulation assays is mainly due to increased cell death in tissue culture. Interestingly, we did not detect increases of Caspase 3 cleavage, indicating that cell death occurs by non-canonical apoptotic pathways. A crucial question is whether lower cellular viability contributes to abnormal human development associated with Down syndrome, including reduced organ size.

The expression of the triplicated chromosome creates an increased metabolic demand for the amino acid serine. As observed in aneuploid yeast cells, we found that human trisomic cells rely on serine to proliferate. Serine is a crucial amino acid that can be converted into glycine, and together, they account for most of the codon usage in the cell. Serine is also used for nucleotide and lipid biosynthesis (Locasale, 2013). We and others previously showed that aneuploid yeasts, trisomic primary fibroblasts, and trisomic MEFs had elevated levels of long-chain bases, which are the direct product of serine palmitoyltransferase (Hwang et al., 2017; Hwang et al., 2019; Tang et al., 2017). Here we found that trisomic human fibroblasts show an increased expression of SPT subunits. Long-chain bases are essential components of the nuclear membrane, and cells harboring an extra chromosome show abnormal nuclear morphologies. A recent study of the metabolome profiles of plasma samples from individuals with Down syndrome revealed serine and sphingosine-1-phosphate, the main form of long-chain-base in the blood, as the top metabolites with lower abundance compared to control individuals (Powers et al., 2019). Importantly, dietary serine can have significant consequences on human physiology (Baksh et al., 2020; Ke et al., 2020). Future research will determine whether a diet enriched with the amino acid serine could have beneficial effects on the health of individuals with Down syndrome.

## Materials and Methods

### Culture of human cell lines

Primary human fibroblasts from euploid individuals and Trisomy fibroblasts (Figure S1A) were purchased from Coriell Cell Repositories. They were maintained in Minimum Essential Media (MEM with NEAA; Gibco 10370088) supplemented with 15 % FBS and 2 mM L-glutamine. To make serine/glycine depleted media, MEM (Gibco 11095080) was supplemented with alanine, asparagine, aspartic acid, glutamic acid, proline and dialyzed serum (3kD MWCO).

### RNASeq

Cells were grown for 48 hours and 2 x 10^5^ cells were harvested between 50 to 70% confluency. The RNAeasy Kit from Qiagen (cat # 74104) was used to purify the RNA and a nanoDrop was used to measure concentration. Samples were shipped for on dried ice for transcriptome sequencing to BGI Americas (https://www.bgi.com/). Paired-end reads were aligned to human primary genome hg38, with star_2.5.3a 1, annotated with GENECODE GRCh38.p12 annotation release 292. Aligned exon fragments with mapping quality higher than 20 were counted toward gene expression with featureCounts_1.5.2 3. Expression normalization was performed using the TPM (Transcripts Per Kilobase Million) method (Dobin et al., 2013; Harrow et al., 2012; Liao et al., 2014). Parameters: Genome: hg38.primary.fa, GTF: gencode.v29.primary_assembly.annotation.gtf.

### RNASeq Data from two other studies

Data from Sullivan et al. study was obtained from the public available database:
https://www.ncbi.nlm.nih.gov/geo/query/acc.cgi?acc=GSE79842 and
https://www.ncbi.nlm.nih.gov/geo/query/acc.cgi?acc=GSE84531. Data from the Letourneau et al. study was obtained from the public available database:
https://www.ncbi.nlm.nih.gov/geo/query/acc.cgi?acc=GSE55426.

### RNASeq analysis

All RNASeq reads were normalized to the total number of reads per experiments (T.P.M. transcripts per million reads). To calculate the fold changes among all cell lines, the average T.P.M per gene was calculated for all the control euploid samples and the average T.P.M was used as a reference genome. Log_2_ (T.P.M. per sample/ T.P.M of reference) was obtained for each sample. We included FC for genes that were detected in all samples and a cutoff of 1 T.P.M. or greater in our analysis. The log_2_ ratios for all the transcriptomes are available in the supplemental tables. Hierarchical clustering was performed using the program Wcluster. WCluster takes both a data table and a weight table to allow individual measurements to be differentially considered by the clustering algorithm. Gene expression data were clustered by a Pearson correlation metric with equal weighting given to all data, or with no weight given to genes on the triplicated chromosomes. The PRISM software was used to calculate frequency distributions, fit to a Gaussian curve, identify of the outliers, perform linear regression analyses, and calculate Pearson r correlation coefficients.

### Western blot assay

Cells were lysed in lysis buffer (25 mM Tris-HCl pH 7.5, 150 mM NaCl, 1% Triton X-100, 0.1% SDS, and 0.5% deoxycholate with protease inhibitors). Lysate was separated on SDS polyacrylamide gels, and then proteins were transferred onto a PVDF membrane and analyzed with antibodies against SPTLC1 (Abcam, ab176706), SPTLC2 (Abcam, ab23696) and GAPDH (Millipore, AB2302). Immunoreactive signals were detected by the Super Signal reagent (Pierce).

### Cell proliferation assay

1 x 10^4^ of cells were plated on 24 well plates with the standard culture media. After 8 hours, cells were washed with PBS and replaced with conditional media with or without serine/glycine (0.4 mM). Cell numbers were counted every 24 hours for 4 days.

### Cell death assay

To distinguish between live and dead cells, cells were stained with LIVE/DEAD™ Fixable Dead Cell stain kit (Invitrogen, L34969). Briefly, 4 x 10^4^ of cells were plated on 60 mm dish. After 2 days, detached cells were incubated with fluorescent reactive dye at room temperature for 30 minutes in dark. Stained cells were washed with PBS and then fixed with PBS with 4 % formaldehyde for 15 minutes. After washing cells with PBS, cells were resuspended in PBS with 1 % bovine serum albumin and analyzed the fixed cell suspension by flow cytometry.

### Protein extraction and digestion

Frozen cell pellets were lysed with RIPA buffer (150 mM NaCl, 1% NP40, sodium deoxycholate 0.5%, SDS 0.1%, TRIS-HCl 25 mM pH 7.4, DTT 5 mM) and insoluble material was removed by pelleting at 20k x g for 10 min. The supernatant was methanol/chloroform precipitated and dried on the benchtop. Dry peptide pellets were resuspended in 8 M urea, 50 mM ammonium bicarbonate (ambic). Protein concentrations were measured using the DC protein assay kit (BioRad). Proteins were reduced by adding DTT to a final concentration of 5 mM and incubating for 30 min at RT, then alkylated with 15 mM iodoacetamide in the dark at RT for 30 min. An additional 5 mM DTT was added to quench the iodoacetamide. The urea concentration was diluted to 2 M by adding 3 volumes of 50 mM ambic. Proteins were digested overnight at room temperature with lysyl endopeptidase (lysC, Wako, Richmond, VA) at a ratio 1:125 enzyme:protein at RT. Samples were diluted to 1 M urea with 50 mM ambic and digested with Trypsin (Promega #V5111) for 10 hours at 1:125 enzyme:protein at 37°C. Digestion was stopped by the addition of formic acid (FA) to a final concentration of 2%. Acidified peptides were loaded onto pre-equilibrated Sep-Pak tC18 cartridges (Waters) and the columns were washed with 1% FA. Bound peptides were eluted with 70% acetonitrile (ACN), 1% FA, dried, and re-suspended in water. Protein concentration was measured using the Pierce colorimetric peptide assay (Thermo, #23275). 50 μg of peptides from each sample was aliquoted and dried down, then re-suspended in 100 ul of 0.5 M HEPES, pH = 8.5. Peptides were labeled by the addition of 0.25 mg TMT in anhydrous ACN for 60 min at RT. Labeling was quenched with 8 μl 5% hydroxylamine and acidified with 16 μl neat FA. A small aliquot (5 μl) of each labeled sample was removed and mixed in equal volumes from each of the 10 channels within each TMT set, desalted on C18 STAGE tips [Rappsilber et al., 2003 - PMID:12585499], and analyzed using a 85 min LC-MS/MS/MS method with synchronous precursor selection (SPS) enabled to generate TMT reporter ions from MS2 fragment ions. Reporter ions were scanned at 60,000 resolution in the orbitrap. The final sample mixes for each 10plex were made based on the relative intensity of total reporter ions from each in order to achieve equal levels.

### Peptide Fractionation

Combined TMT-labeled peptides were de-salted, resuspended in 300 μl buffer A (5% ACN, 10 mM NH_4_HCO_3_, pH 8) and separated by high-pH reverse-phase HPLC (PMID: 21500348, Wang et al. Proteomics 2011) on a C18 column (Waters, #186003570, 4.6 mm x 250 mm, 3.5 μ ID) using a 50 min gradient from 18% to 38% buffer B (90% ACN, 10 mM NH_4_HCO_3_, pH 8) with a flow rate = 0.8 ml/min.. Fractions were collected over 45 min at 28 sec intervals beginning 5 min after the start of gradient in a 96-well plate. These original fractions were pooled into 12 samples each containing four fractions (only 48 of 96 fractions were used) by sampling at equal intervals across the gradient, i.e. by combining fractions 1/25/49/73, 3/27/51/75, 5/29/53/77, ....13,37,61,95. This pooling strategy minimizes overlap between fractions. The pooled samples were dried down, resuspended in 50 μl of 5% FA, and desalted on STAGE tips.

### LC-MS/MS Analysis

Each fraction was analyzed on a Thermo Orbitrap Fusion mass spectrometer (Thermo Fisher Scientific) equipped with an Easy nLC-1000 UHPLC (Thermo Fisher Scientific). Peptides were separated with a gradient of 6–25% ACN in 0.1% FA over 115 min and introduced into the mass spectrometer by nano-electrospray as they eluted off a self-packed 40 cm, 75 μm (ID) reverse-phase column packed with 1.8 μm, C18 resin (Sepax Technologies, Newark, DE). They were detected using the real-time search (RTS) MS3 method [PMID: 30658528] which employs a data-dependent Top10-MS2 method and a RTS triggered SPS-MS3 to collect reporter ions. For each cycle, one full MS scan was acquired in the Orbitrap at a resolution of 120,000 with automatic gain control (AGC) target of 5 × 10^5^ and maximum ion accumulation time of 100 ms. Each full scan was followed by the individual selection of the most intense ions, using a 2 sec cycle time for collision-induced dissociation (CID) and MS2 analysis in the linear ion trap for peptide identification using an AGC target of 1.5×10^4^ and a maximum ion accumulation time of 50 ms. Ions selected for MS2 analysis were excluded from reanalysis for 60 s. Ions with +1, >+5, or unassigned charge were excluded from selection. Collected MS2 spectra were searched in real time against a database of all uniprot reviewed human sequences (uniprot.org, downloaded May 1, 2019) using binomial score cutoff of 0.75. Passing spectra triggered synchronous precursor selection of up to 10 MS2 ions for HCD fragmentation and MS3 scanning in the orbitrap at resolution = 60,000 using a maximum ion time of 200 ms. The gene close-out feature was enabled and applied to each set of 12 fractions and used to limit the selection and collection of MS3 reporter ions to a maximum of 10 peptides per protein across all 12 runs in each 10plex set.

### Peptide Searching and Filtering and Protein Quantification

MS/MS spectra were then searched to match peptide sequences using SEQUEST v.28 (rev.13) (Eng et al., 1994) and a composite database containing all reviewed human uniprot protein sequences (uniprot.org, downloaded May 1, 2019) and their reversed complement. Search parameters allowed for two missed cleavages, a mass tolerance of 20 ppm, a static modification of 57.02146 Da (carboxyamidomethylation) on cysteine, and dynamic modifications of 15.99491 Da (oxidation) on methionine and 229.16293 Da on peptide amino termini and lysines.

Peptide spectral matches were filtered to a 2% false-positive rate using the target-decoy strategy [PMID: 17327847] combined with linear discriminant analysis (LDA) [PMID: 21183079] using the SEQUEST Xcorr and ΔCn scores, mass error, charge, and the number of missed cleavages. Further filtering based on the quality of quantitative measurements (reporter ion sum signal-to-noise ≥ 200, isolation specificity >0.7) resulted in a final protein FDR < 1% for both 10plex experiments.

Total protein intensity values for each sample were derived from the summed reporter ion intensities from the corresponding channel from all peptides mapping to that protein. Values were normalized to account for small variations in sample mixing based on the sum intensity of values from all proteins in each channel within each 10plex. Relative abundances for each protein across samples within each 10plex were calculated as the fraction of the total intensity derived from each channel. The average log_2_[Trisomy21/WT] was calculated and used for all subsequent analysis.

## Acknowledgements

This research was supported by a grant from the National Institutes of Health grant 1R01GM118481-01A1 to EMT.

## Supplemental Figure Legends

**Figure S1.**
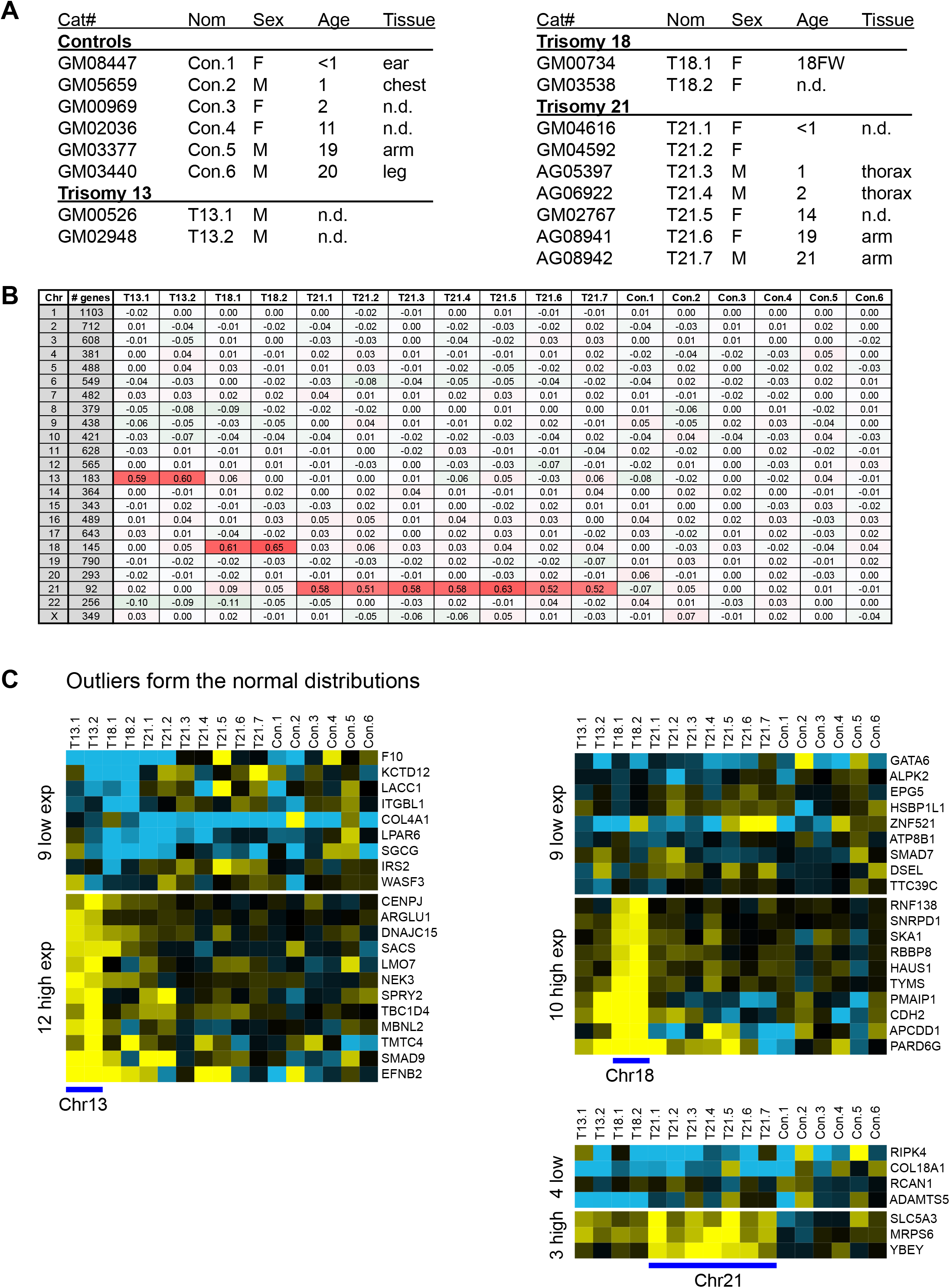
Transcript levels increase proportionally with gene copy number in trisomic primary fibroblasts. Cell lines utilized in this study. All cell lines were obtained from the Coriell Institute (https://www.coriell.org). Cat# = Catalogue number, Nom = nomenclature used in the manuscript. Sex, age, and tissue of the donor are listed. **B.** The values of the averaged gene expression per chromosome calculated for each cell line are shown. The triplicated chromosomes are highlighted in red. **C.** Gene expression of triplicated genes that are outliers in the fits of the normal distributions for each trisomic cell lines is shown. Outliers were identified by fitting the distributions to a Gaussian curve using least squares regression methods and defining the points above or below the curve.

**Figure S2.**
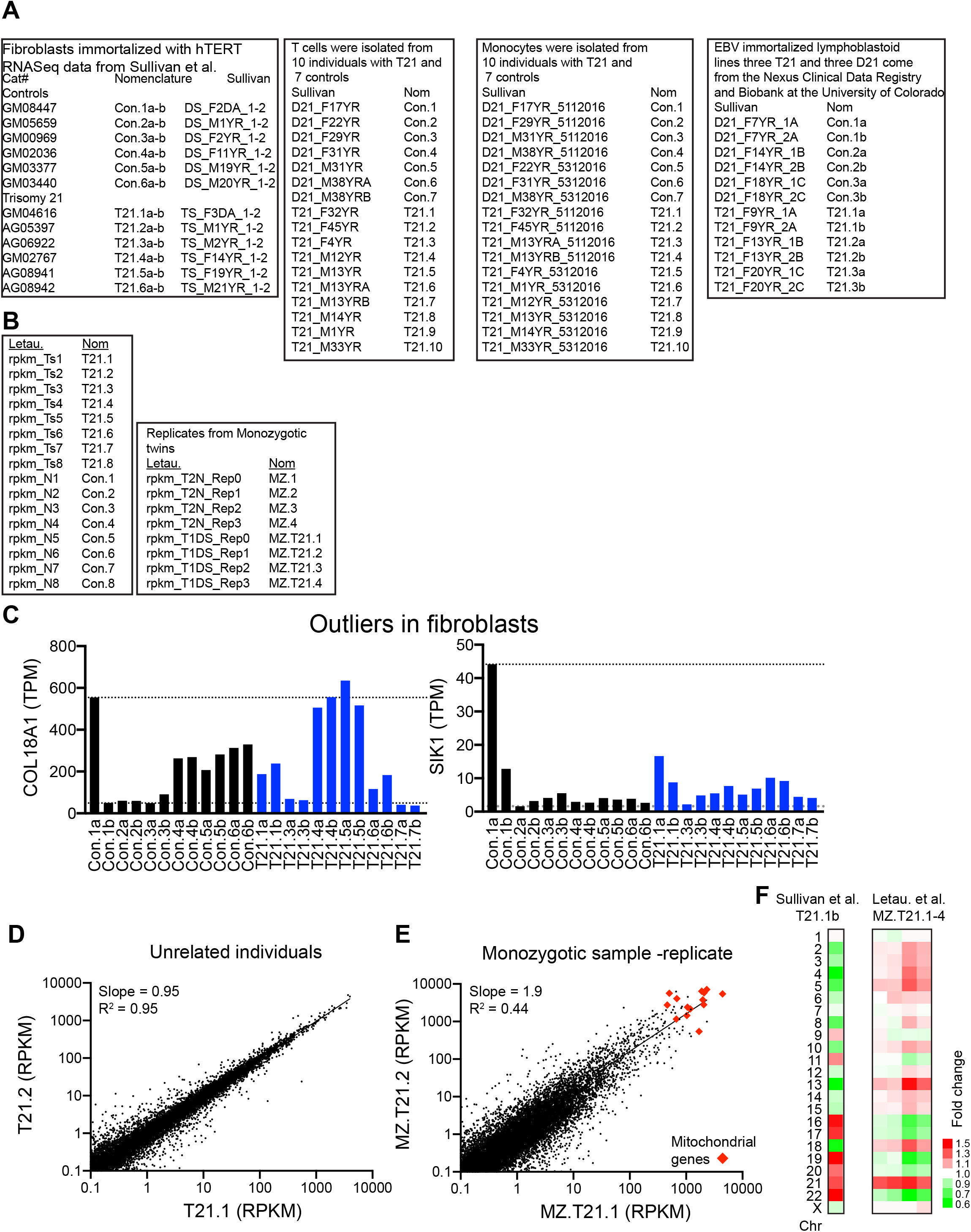
Transcript levels increase proportionally with gene copy number in distinct trisomic cell lines. **A.** Nomenclature of the cell lines analyzed by Sullivan et al. **B.** Nomenclature of the cell lines analyzed by Letourneau et al. **C.** Example of a couple of outliers of the fits of the distributions of triplicated genes in the data for the fibroblasts analyzed by Sullivan et al. **D.** Linear regression analysis of the RNA counts of two unrelated fibroblasts transcriptomes, and, (**E)** the monozygotic twins analyzed by Letourneau et al. **F.** Comparison of the average gene expression per chromosome of the outlier transcriptomes in the Sullivan et al. and Letourneau et al. studies.

**Figure S3.**
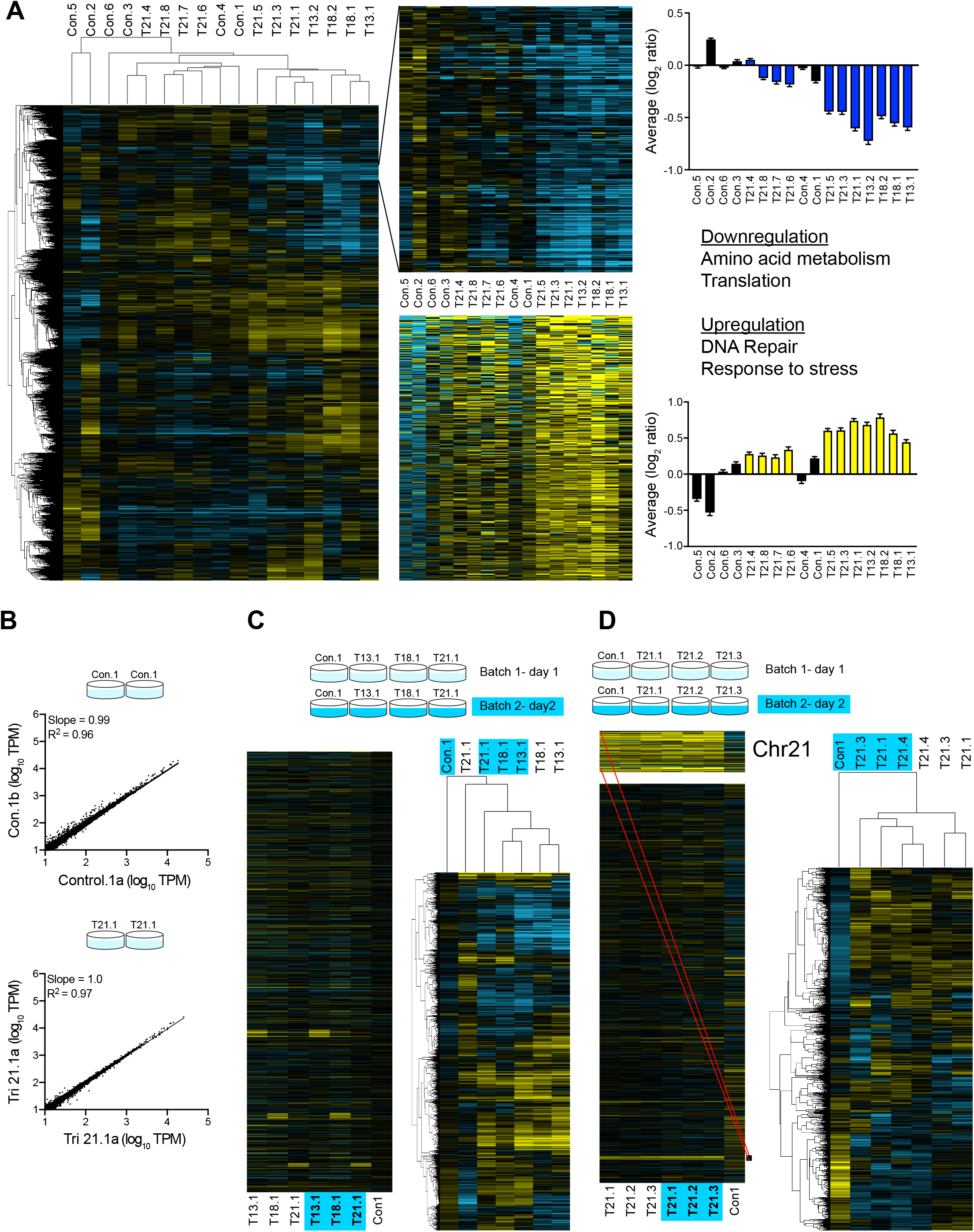
Gene expression patterns of human primary fibroblasts. **A.** Hierarchical clustering analyses of the expression patterns of primary fibroblasts analyzed in this study (left heatmap). Down (middle top) and upregulated (middle bottom) cluster of genes in primary trisomic fibroblasts. Bars represent the average change in gene expression of the down and unregulated clusters (right). **B.** Linear regression analysis of the transcriptome profiles of one control and one trisomy 21 cell lines grown in parallel show a high degree of reproducibility. **C-D**. Hierarchical clustering analyses of the expression patterns of primary fibroblasts grown months apart show that culture conditions significantly influence the patterns of gene expression. Cluster patterns are driven by the time of culture in each comparison (blue boxes).

**Figure S4.**
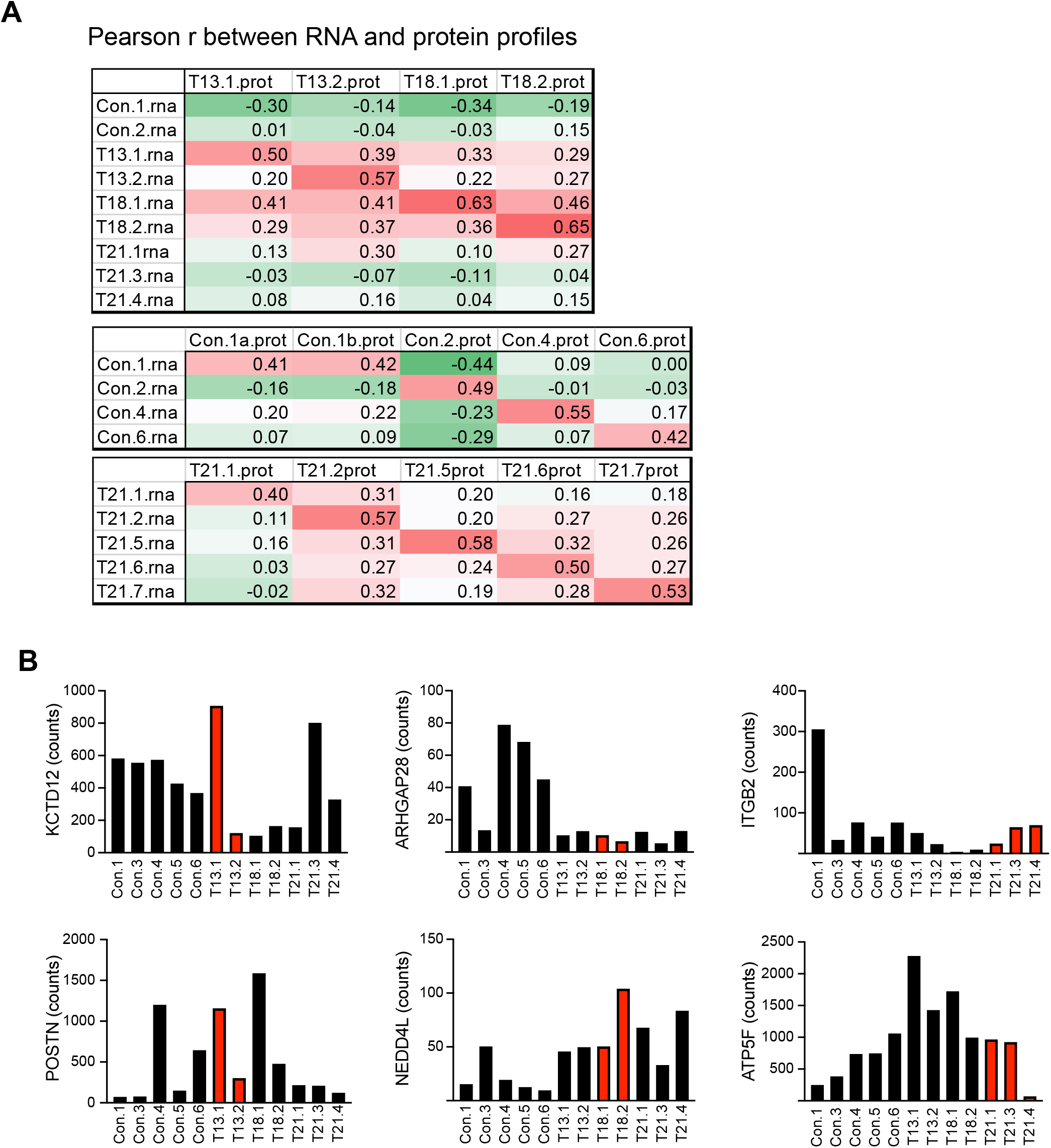
Protein levels proportionally increase with copy number in trisomic fibroblasts. **A.** Pearson correlation coefficients were calculated for the transcriptome and proteome profiles of the cell lines indicated. **B.** Examples of the peptide counts of a few triplicated genes that are outliers in the fits of the normal distributions. Red bars correspond to peptide counts of the triplicates genes.

**Figure S5.**
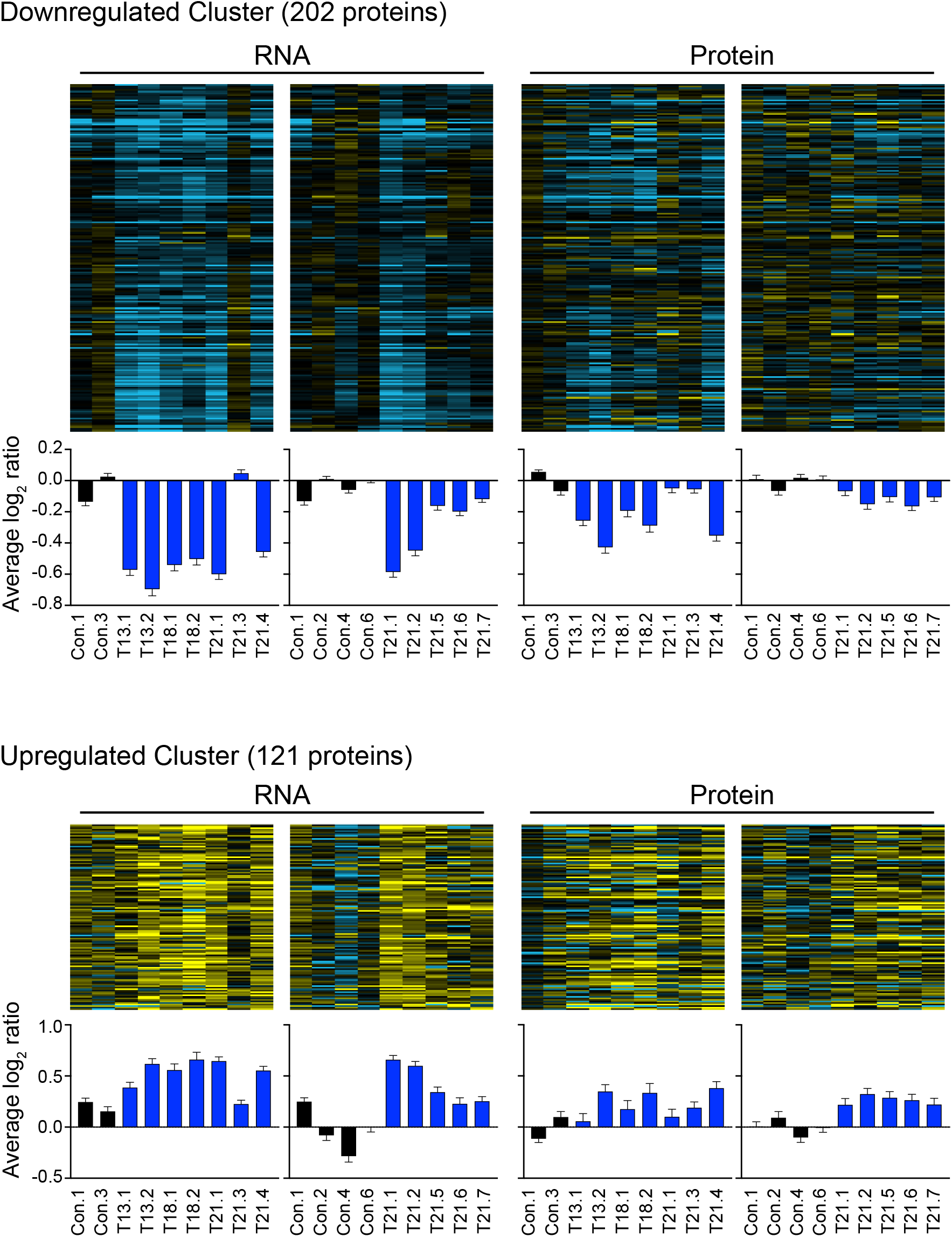
Cluster of down and upregulated genes in trisomic fibroblasts. Down and upregulated clusters of genes identified in Figure S3 lead to changes in protein levels. Bars represent average change in transcript (left) and protein levels (left) in each of the cell lines analyzed.

**Figure S6.**
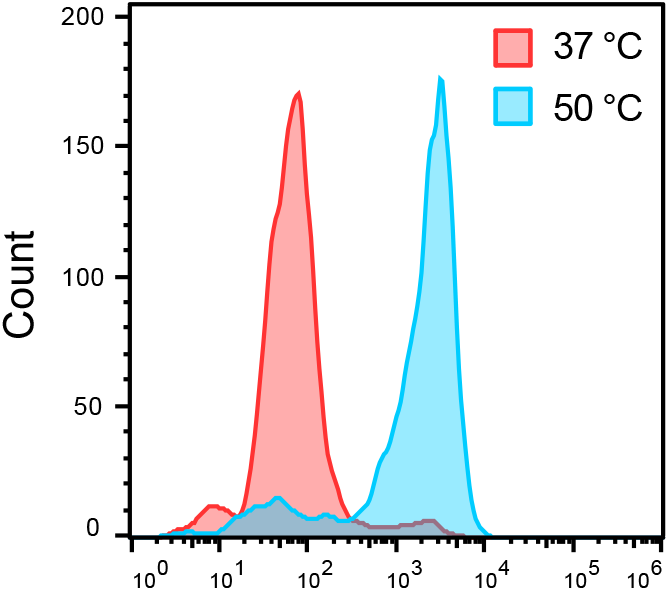
Assay for cell viability. The LIVE/DEAD™ Fixable Green Dead Cell Stain Kit uses a dye that reacts with free amines yielding intense fluorescent staining. In viable cells (red), the dye’s reactivity is restricted to the cell surface resulting in less intense fluorescence than non-viable cells (blue) wherein the dye diffuses inside the cell.

